# Early development of Neanderthals revealed through virtual microanatomy

**DOI:** 10.64898/2026.02.25.707915

**Authors:** Justyna J. Miszkiewicz, Ricardo Miguel Godinho, Anne Marie Sohler-Snoddy, Kerstin Pasda, Florent Détroit, Patrick Mahoney, Thomas Rathgeber, Cosimo Posth, Thorsten Uthmeier, Alvise Barbieri

## Abstract

The ontogeny of Neanderthal (*Homo neanderthalensis*) perinates is poorly understood due to the paucity of juvenile skeletal remains. Here we reconstruct fetal bone growth, and explore deciduous tooth structures, in three Neanderthal juveniles (Sesselfelsgrotte, 1, 2 and 3) (90,000–50,000 years ago) from southeastern Germany using non-destructive microcomputed tomography.

Sesselfelsgrotte 1 exhibited bone tissue consistent with modern human perinatal plexiform-like structures and primary osteons. Long bones showed regions of advanced growth compared to the mandible and frontal bone, which can be explained through different processes of ossification and potentially localized faster development in Neanderthals compared to modern humans. Bone microstructure resembles that of the late third trimester of modern humans, agreeing with previous estimates based on macroscopic data. Sesselfelsgrotte 2 and 3 deciduous teeth retain hypodensities deep within the crown dentine consistent with interglobular dentine.

We conclude that the fetal bone patterning is similar to modern humans with areas of advanced growth, indicating that the growth trajectory for this Neanderthal perinate was broadly equivalent to that of modern humans. The abnormal dentine mineralization points toward a possible systemic disorder.

## 1. Introduction

Neanderthals (*Homo neanderthalensis*) have long captivated our curiosity because they co-existed with modern humans for at least 5k years (Mylopotamitaki et al., 2024) and are relatively well represented in the fossil record (Guil-Guerrero & Manzano-Agugliaro, 2023). Their skeletal and dental remains have allowed palaeoanthropologists to reconstruct aspects of Neanderthal lives including lifestyle, culture, diet, environmental adaptation, and their genome has revealed inbreeding with modern humans (Guil-Guerrero & Manzano-Agugliaro, 2023; Petr et al., 2020; Stringer, 2025; Stringer & Crété, 2022). The Neanderthal fossil record, however, includes limited infant remains (Golovanova et al., 1999; Maureille, 2002b, 2005; Peyrony, 1930; Rak et al., 994; Rathgeber, 2006), some of which have been used to establish biological similarities and differences with modern human development and the juvenile period trajectories (Bastir et al., 2007; García-Martínez et al., 2020; Gunz et al., 2012; Ponce de León et al., 2008).

Sesselfelsgrotte (Essing, Bavaria) is one of the richest and most unique Neanderthal sites in western Europe, preserving bones and teeth of three non-adult individuals (Sesselfelsgrotte 1 to 3) indirectly dated to between 90 and 50 ka (Freund, 1998; Rathgeber, 2006; Richter et al., 2000). These remains consist of one deciduous upper left second molar (Sesselfelsgrotte 2), one possible deciduous lower left second molar (Sesselfelsgrotte 3), and 12 bone fragments previously ascribed to a perinate (Sesselfelsgrotte 1) based on taxonomic determination and external bone measurements performed two decades ago (Rathgeber, 2006). In this study, we take advantage of these rare osteological and dental specimens to expand the current knowledge on Neanderthal ontogeny in relation to modern humans. We perform a comparative virtual microanatomy study using microcomputed tomography (micro-CT) scans of the skeletal remains of the Sesselfelsgrotte 1 to 3 individuals. Previous studies on immature Neanderthals have demonstrated the usefulness of this virtual approach to assess skeletal microstructure, which retains evidence of living cell activity linked to the formation and growth of bone and tooth tissues (e.g., Colombo et al., 2019; Mahoney et al., 2021; Nava et al., 2020; Sawada et al., 2004).

We use virtual microanatomy to describe bone formation processes for Sesselfelsgrotte 1. Comparisons are undertaken with previously published data for modern humans and other Neanderthals, including previous micro-CT scans of La Ferrassie 4bis and Le Moustier 2 Neanderthal infants. The two main aims of our study are to (i) assess whether bone microanatomy corresponds with the proposed fetal third trimester stage of development for Sesselfelsgrotte 1 and in doing so expand current knowledge of ontogenetic bone formation in Neanderthal foetuses and infants, and (ii) explore the internal microstructure of Sesselfelsgrotte 2 and 3 deciduous molars.

### 1.1. Bone growth and microstructure in modern human foetuses and infants ≤ 2 years old

Human skeletal development from the fetal to birth stages is an intricate biological process (e.g. Cunningham et al., 2016; Gardner & Gray, 1970; Land & Schoenau, 2008; Nemec et al., 2011; Salle et al., 2002; Scheuer & Black, 2004; Thomson, 1951; Weaver & Fuchs, 2014). In this section, we summarise this development with the goal of introducing key terms that will be used in our study. We will first provide an overview of skeletal development and growth processes, followed by bone histology and microanatomy changes through these developmental milestones. We will follow the terminology outlined by Moreno et al. (2025), referring to *in utero* stages as ‘prenatal’ or ‘fetal’, and ‘perinatal’ when in reference to up to two months before birth, although this also includes up to seven days postnatally.

The human skeleton begins formation *in utero* and starts to ossify through intramembranous and endochondral processes between the 7^th^ to 8^th^ week of the first trimester (Johansen et al., 2007). Mesenchymal cells differentiate into bone forming osteoblasts and ossification occurs on a fetal skeletal scaffold made up of fibrous membranes and hyaline cartilage (Cunningham et al., 2016). Around 8 weeks *in utero*, somites (mosoderm tissue that gives rise to parts of the musculoskeletal system) disappear, and joint formation begins (Chamley, 2005). Ossification occurs from within primary centres on the post-cranial cartilaginous scaffold, leading to the creation of long bone shafts. This process is known as endochondral ossification (Cunningham et al., 2016). Most of the skull bones form through intramembranous ossification where bone tissue is deposited directly without a preexisting cartilaginous model (White & Wallis, 2001). As the baby grows over the three trimesters of pregnancy, bone tissue continues forming, resulting in about 275 bones at birth. Shafts of long bones, such as the femur, lengthen from about 34 mm at 16 weeks to 95.5 mm at 41 weeks (Salle et al., 2002). By 20 weeks, the bone matrix is highly organized in terms of its mineralization (Glorieux et al., 1991). Until birth, fetal and perinatal bone microstructure receives great blood supply, vascularising bone tissue. The shafts expand in diameter, while they ossify and are modelled (Salle et al., 2002) and remodelled partly with mechanical stimulation from fetal movement and kicks (Burton et al., 2008; Verbruggen et al., 2018). This process is executed by osteoblasts and osteoclasts which form and resorb bone tissue either independently, changing the shape of bone (modelling), or teamed up, replacing old bone with new tissue (remodelling). Histology from modern human foetuses aged 5 months *in utero* (20 weeks) shows the existence of osteocytes (mature osteoblasts trapped in calcified bone matrix), osteoblasts, and osteoclasts, indicating that both remodelling and modelling start early *in utero* (Ernst et al., 2011).

The extensive blood supply during rapid growth results in expansive vascularity seen in fetal bone microstructure (Filipowska et al., 2017). In the first trimester, the main bone tissue type present is woven, which has a haphazard collagen orientation owing to its rapid deposition (Shapiro & Wu, 2019). In the second trimester, woven bone still dominates in a cross-section of a long bone, but it also starts to incorporate small amounts of primary lamellar bone that appear parallel-fibered (Figure 1a–c) (Shapiro & Wu, 2019). Long bone cross-sections at this stage appear trabeculae-like, with elongated regions of bone enveloping the medullary cavity separated by large spaces (Figure 1d) (Dhawan et al., 2014), increasing structural integrity well into the third trimester. Various anatomical studies of individual modern human bones from *in utero* autopsies have evidenced this micromorphology, including the shaft of the tibia at 4-4,5 months (Shapiro & Wu, 2019); the femur at 5 months (21+5 weeks) (Figure 1d, Dhawan et al. 2014: 276), the rib, tibia, and humerus at 5-8 months (22–32 weeks) (Chu et al., 2024; Eleazer, 2007).

**Figure 1.**
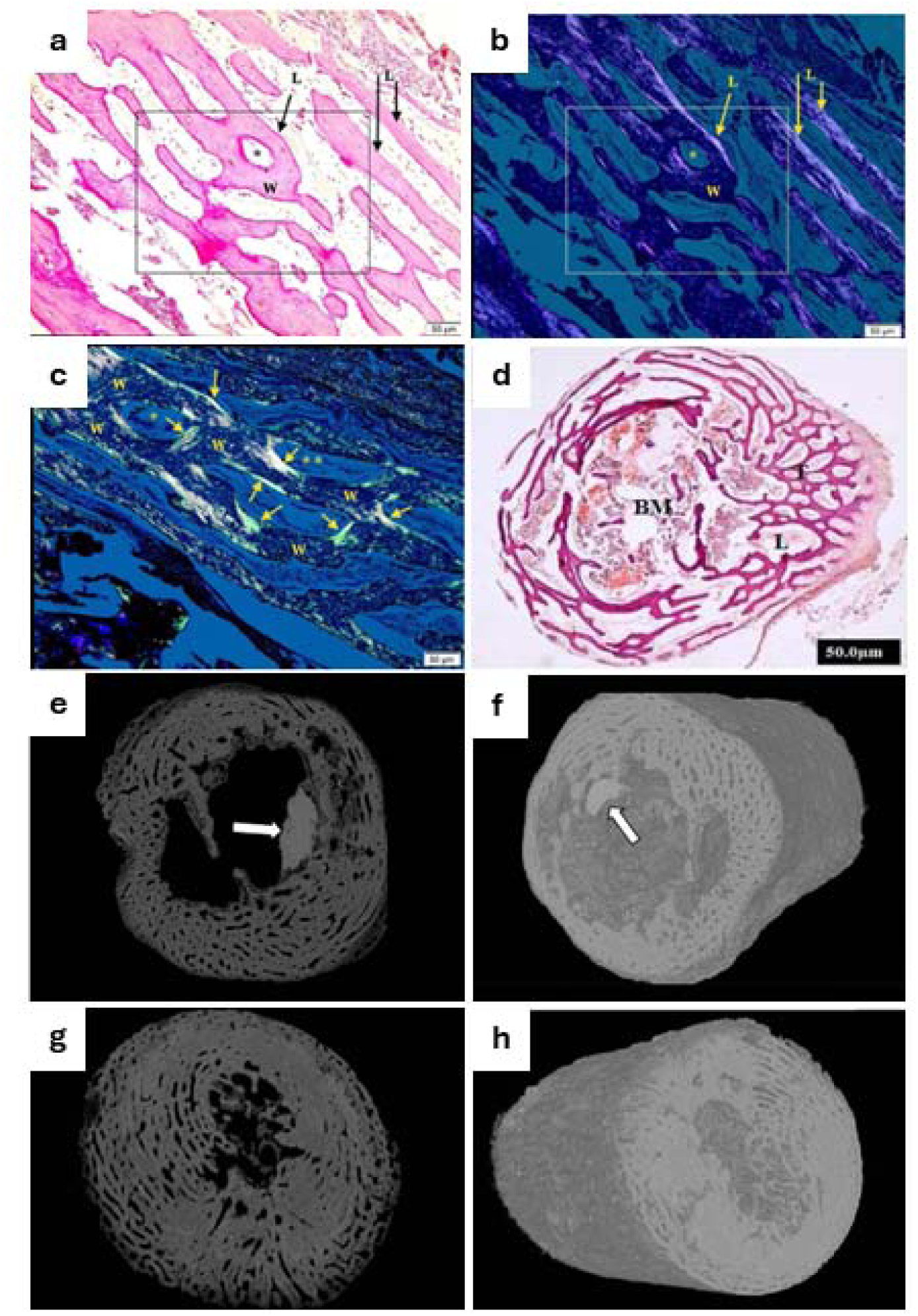
Histology (a–d) and microCT (e–h) images showing modern human bone microstructure in long bone shafts from second and third trimester stages of development. Images a–c are from 4–4,5 months (second trimester) tibial cortices showing trabecular-like structure of cortical bone (a–c), and presence of a combination of woven (W) and lamellar (L) bone (indicated with arrows). Scale bar in a–c is 50μm. Image d shows histology in a 5 month (21+5 weeks, second trimester) femoral diaphysis illustrating trabecular structure of cortical bone filled with bone marrow (BM). Images e–f are from the collection of St. Mary Spital (UK) showing a diaphysis of the left femur from 8 months (30–32 weeks, third trimester) showing continued trabecular, plexiform-like structure of cortical bone with increasing integrity and density. Images g–h are from the Stothard Place collection in 8 months (30–32 weeks, third trimester), showing denser trabecular structure with plexiform-like composition. No scale bars were provided with the original images. The white arrow in e–f is from the original publication indicating concentration of bacterial bioerosion. Images a–c are reprinted based on SA 4.0 open access license from Shapiro & Wu, 2019, p. 149. Image d is reprinted from Dhawan et al. 2014, p. 276 with permission from the European Journal of Anatomy (17 July 2025). Images e–h are reprinted from Booth et al. 2016, p. 130–131 with permission from Elsevier under license number 6160190936241.

Approaching the perinatal stage, primary lamellar bone, which is more organized than woven, and arranged in a characteristic manner with layers of superimposed bone, develops along with primary osteons which are layers of lamellar bone enveloping longitudinal blood canals (Shapiro & Wu, 2019). This bone tissue is still combined with woven bone, forming a characteristic appearance that is variably referred to as ‘plexiform’ or ‘parallel-fibred’ bone (Figure 1e–h) (Caccia et al., 2016; Goldman et al., 2009; Pfeiffer, 2006; Shapiro & Wu, 2019). This type of bone has also been observed in modern human infants, children, and teenagers, during growth spurts (Goldman et al., 2009; Pfeiffer, 2006). The term “plexiform” has introduced some confusion in the literature as this type of bone is not true plexiform as that found in other mammals where it has a distinct brick wall-like appearance (Goldman et al., 2009; Pfeiffer, 2006).

Postnatally, and until two years old, woven bone is no longer predominant as it is replaced with tissues that have better organized collagen fibres. This bone tissue type is still primary but has a lamellar structure that contains primary osteons (Pfeiffer, 2006; Sawada et al., 2004). Very little regions of woven bone are left by two years old (Pfeiffer, 2006). Isolated secondary osteons will start emerging and progressively increase in density well into adulthood. Sporadic secondary osteons were observed in those aged >1 years in modern human ribs and femora (Pfeiffer, 2006; Sawada et al., 2004). In a recent study examining humeral microstructure prenatally to 1.5 years, Moreno et al. (2025) noted the presence of secondary osteons at 6 months, but not before. An earlier study by Baltadjiev (1995) discussed the existence of ‘osteons’ and ‘Haversian canals’ in the tibiae of modern human foetuses, but it was referring to primary rather than secondary osteons since no cement lines were observed. Pfeiffer (2006) noted possible presence of regions of “plexiform” bone in the ribs from archaeological 0–2-year-olds. One–two-year-olds have infrequent circumferential lamellae, which become better established after 3 years old (Sawada et al., 2004).

### 1.2. Modern human dental growth and development

Odontogenesis is a complex process that starts as early as six weeks after fertilization with the formation of the dental laminae (Schoenwolf et al., 2014). Deciduous tooth formation ensues in the central incisors at about 14–16 weeks after conception (Hillson, 1996). By birth, all deciduous tooth types have started mineralization, and all deciduous teeth have fully erupted by approximately ∼2.5 years old post birth (AlQahtani et al., 2010; Scheuer & Black, 2000). Root lengths vary as the crown erupts but are typically half the length of the crown at the time of tooth eruption (Liversidge & Molleson, 2018).

Dentine forms the bulk of the tooth, and it is covered by enamel on the crown and by cement on the root (Ireland, 2010). During dentine formation, which begins around 14 weeks of development, odontoblasts migrate apically (i.e., towards the future tip of the root) away from the basement membrane (which originates the enamel dentine junction), depositing an organic extracellular matrix known as predentine (Ireland, 2010). Mineralization proceeds below the predentine matrix forming front, through seeding of hydroxyapatite crystals in the predentine matrix, forming primary dentine, which is about 70% inorganic by weight (Ireland, 2010). The seeded apatite crystals grow radially, forming spherical bodies (calcospherites) that typically coalesce with other crystals, mineralizing the dentine. Occasionally, these calcospherites fail to fuse resulting in irregular spaces of unmineralized dentine (also known as interglobular dentine, IGD) (Hillson, 1996). Although small amounts of IGD are a clinically common finding, more extreme expressions of this defect may indicate a systemic disorder of mineral metabolism during growth such as rickets (Abe et al., 1988; Seow et al., 1989).

Amelogenesis (i.e., enamel formation) is initiated almost immediately after dentinogenesis has started (Schoenwolf et al., 2014). In the secretory phase, a matrix is deposited by ameloblasts that include an organic (primarily amelogenin preoteins, but also enamelin and ameloblastin) and an inorganic component (hydroxyapatite crystals). Initial enamel deposition is only partially mineralized and is ensued by maturation, which involves the removal of organic material and water by the ameloblasts and the influx of calcium and phosphate ions (Hillson, 1996). The hydroxyapatite crystals grow, and fully matured enamel consists of ∼97% inorganic material by weight (Ireland, 2010). It is acellular, making it incapable of repair or regeneration following eruption.

Once the tooth crown is complete, tooth growth proceeds into the root, which is composed of dentine coated by cementum, a partially mineralized connective tissue formed by cementoblasts (Schoenwolf et al., 2014). Root odontoblasts differentiate from dental papilla cells in a process induced by the Hertwig’s epithelial root sheath, which comprises both internal and external enamel epithelium (Harris, 2015). Together, these sequential and spatially coordinated processes produce the mature tooth, with distinct hard tissues exhibiting specialized histological and functional properties.

### 1.3. Previous fossil record for Neanderthals <2 years old

The fossil record for immature Neanderthals (< 2 years old) is incredibly scarce. The following neonates and infants are known: 2-week-old Mezmaiskaya 1 and 1 to 2-year-old Mezmaiskaya 2 from Russia (Golovanova et al., 1999); <120 days old Le Moustier 2 from France (Maureille, 2002b, 2005; Peyrony, 1930); 1.5–1.7-year-old Dederiyeh 1 from Syria (Sasaki, 2003); 10-month old Amud 7 from Israel (Rak et al., 1994); ∼12-days or 6,5 weeks old La Ferrassie 4bis (see details below), and 22 to 24-month-old Ferrassie 8 from France (Chevalier et al., 2021; Heim, 1976, 1982a; Maureille, 2002a). Only two potential Neanderthal foetuses have been so far discovered: the 8 to 9-month-old Sesselfelsgrotte 1 from Germany (Rathgeber, 2006); and 7-month-old La Ferrassie 5 from France (Heim, 1976, 1982a). Although La Ferrassie 4 was originally aged as a 8.5-month-old foetus (Heim, 1982b) it was later identified in fact as being part of Le Moustier 2 (Maureille, 2002a, 2002b) and so it is excluded as one of the existing potential foetuses of the Neanderthal record. Ontogenetic research based on some of these specimens, while limited, has revealed that, for example, deep and short ribcages of Neanderthals were present at birth (García-Martínez et al., 2020), their brain size at birth was likely similar to *Homo sapiens* and with comparable obstetric constraints (Gunz et al., 2012; Ponce de León et al., 2008); but that their spatio-temporal processes underlying craniofacial transformations during early ontogeny differed from modern humans (Bastir et al., 2007; Freidline et al., 2013). Further research supports that such differences in facial morphology result from contrasting growth and development processes impacting modelling and/or remodelling (Lacruz et al., 2015). In the post-cranial skeleton, Sawada et al. (2004) examined the histology of the femur midshaft cross-section in the 1.5–1.7-year-old Dederiyeh 1 Neanderthal child to find that its cortical thickness matched that of 5–6-year-old modern human (i.e., *Homo sapiens*) children. Secondary osteon presence and formation patterns were similar to modern human children older than 2 years old (Sawada et al., 2004). On this basis, Sawada et al. (2004) noted that bone development in Neanderthals might have been more advanced than in modern human children. However, the histomorphology of primary bone in the Dederiyeh 1 specimen was similar to modern human children aged 1–2 years. They concluded that Neanderthal bone growth in the early stages of ontogeny was likely different from modern humans since they observed a combination of bone tissues that matched modern human children at different ages. The perinatal bones of the Sesselfelsgrotte 1 specimen from Germany were examined by Rathgeber (2006) who estimated that they were 8 months old based on comparisons with modern human and the La Ferrassie foetuses (although the bones of La Ferrassie 4 have now been reconciled with Le Moustier 2). This age estimate is on the cusp of potential birth presenting a unique opportunity to evaluate bone microstructure and Neanderthal early life ontogeny in this perinate. No study has yet examined bone microstructure in such young Neanderthal fossils, with our current knowledge in this area stemming only from bone histomorphology of the Dederiyeh Neanderthal child (Sawada et al., 2004).

Previous studies of dental development that counted the number of perikymata reported that development was either comparable (Guatelli-Steinberg et al., 2005) or was advanced in (Ramirez Rozzi & Bermudez de Castro, 2004) the teeth of Neanderthals compared to modern humans. A similar situation has been reported in histological studies that examined dental microstructures in permanent teeth, reporting that enamel growth in Neanderthals could be faster or similar to modern humans (Macchiarelli et al., 2006; Smith et al., 2010; Smith et al., 2007). Virtual microanatomical examination of deciduous teeth reported faster growth and a shorter period of development for Neanderthal incisors, canines and molars, when compared to modern humans (Mahoney et al., 2021).

Dental growth and development may be disturbed by various factors, which, when sufficiently severe, may be macroscopically visible on tooth surfaces in the form of enamel hypoplasia (i.e., enamel surface defects such as pits, furrows and bands) (Collignon et al., 2022; Goodman & Rose, 1990). High frequencies of dental enamel hypoplasia have been reported in Neanderthals, in some cases clustering within the range of modern human groups with high incidences of such defects (Guatelli-Steinberg et al., 2004; Hutchinson et al., 1997; Molnar & Molnar, 1985; Ogilvie et al., 1989). A more recent study further examining such developmental disorders found comparable disturbance severity between Neanderthals and other modern human groups, despite Neanderthals having shallower hypoplasia (possibly due to faster tooth development) (McGrath et al., 2021). Moreover, despite Neanderthals and Upper Palaeolithic modern humans showing apparent comparable probabilities of developing hypoplasia, Neanderthals seem to form them at later ages than modern humans (Guatelli-Steinberg et al., 2004).

Other studies examining tooth developmental defects at a microscopic level report dentin mineralization defects ( Smith et al., 2009; Sognnaes, 1956). While the early report of Sognnaes (1956) related those inter-globular dentin defects to vitamin D deficiency, it has also been hypothesized they may be of developmental rather than pathological origin (D’Ortenzio et al., 2018).

Considering the limited number of Neanderthal skeletal and dental elements investigated with microanatomical methods, the remains from Sesselfelsgrotte offer a rare opportunity to further our knowledge on Neanderthal ontogeny and development.

## 2. Materials and Methods

### 2.1. Skeletal and dental remains and archaeological context

The Neanderthal bones and teeth in our study originate from the archaeological site of Sesselfelsgrotte (Essing, Kelheim, Bavaria). These fossils are permanently stored and curated at the Prehistoric and Protohistoric Museum of the UFG Institutes based at the Friedrich-Alexander-Universität (FAU) in Erlangen (Bavaria). Sesselfelsgrotte is a rockshelter located in the lower Altmühl Valley (Essing near Kelheim, Southeastern Germany), a region rich in Palaeolithic sites (Barbieri et al., 2022; Birkner, 1936; Böhner, 2008; Freund, 1998;

Obermaier & Wernert, 1914; Street et al., 2006) (Figure 2). Previous excavations at the shelter unearthed a 7 m deep sequence of limestone eboulis, in which Middle, Upper, and Late Palaeolithic stays as well as historical occupations were buried (Freund, 1998). The Middle Palaeolithic remains include a large faunal assemblage, microfauna remains, fire residues – including potential hearths (Freund, 1998), and a total of 14 human skeletal remains. These consist of the 12 potential fetal bones known as Sesselfelsgrotte 1 (skeletal elements are detailed in Table 1), the fragmented buccal half of an upper left deciduous second molar labelled as Sesselfelsgrotte 2, and the fragmented approximately lingual half of a tentatively identified inferior left deciduous second molar known as Sesselfelsgrotte 3 (Rathgeber, 2006) (Figs. 4 and 5). Sesselfelsgrotte 2 was identified during the excavation of the site in 1968–70s (Freund, 1998). It was uncovered in layer M2, which exhibits a weighted thermoluminescence age of 75.9 ± 3 ka based on five dated samples (recalculated by us from original 73.2±11.7ka in Richter [2001: 37, 39] based on excluded samples that were not indicated of Neanderthal occupation) (Richter, 2001). Contrarily, Sesselfelsgrotte 1 and 3 were identified only in the 1990s, long after their excavation (Rathgeber, 2006). Sesselfelsgrotte 3 was unearthed from layer G2, while apart from the femur and the fibula, all the skeletal remains of Sesselfelsgrotte 1 were uncovered from layer G5 in square B7. The two long bone fragments were uncovered from a collapse of the west excavation profiles of squares B7 – Z7, where the G layers were exposed (Freund 1998, 46). Since their identification, these fetal remains have been interpreted as belonging to the same individual and from the same deposit, G5 (Rathgeber, 2006). Thermal luminescence ages from layer G4 in square A7 date the 15 cm-thick sediment covering Sesselfelsgrotte 1 to 51.1 ± 10.3 ka and 57.5 ± 12.8 ka (also recalculated by us from original 56.0±1.9ka in Richter [2001: 39]) (Richter, 2001). As the deposits between G4 and M1 are undated, we conclude that the Neanderthal fetus (located in layer G5), dates either to the very end of MIS 4 or – given the fact that the thickness of the entire G-Layers-Complex measures 0.5 m only and is underlain by layers I to L with a highly arctic small mammal fauna – more probably to the very beginning of MIS 3.

**Figure 2.**
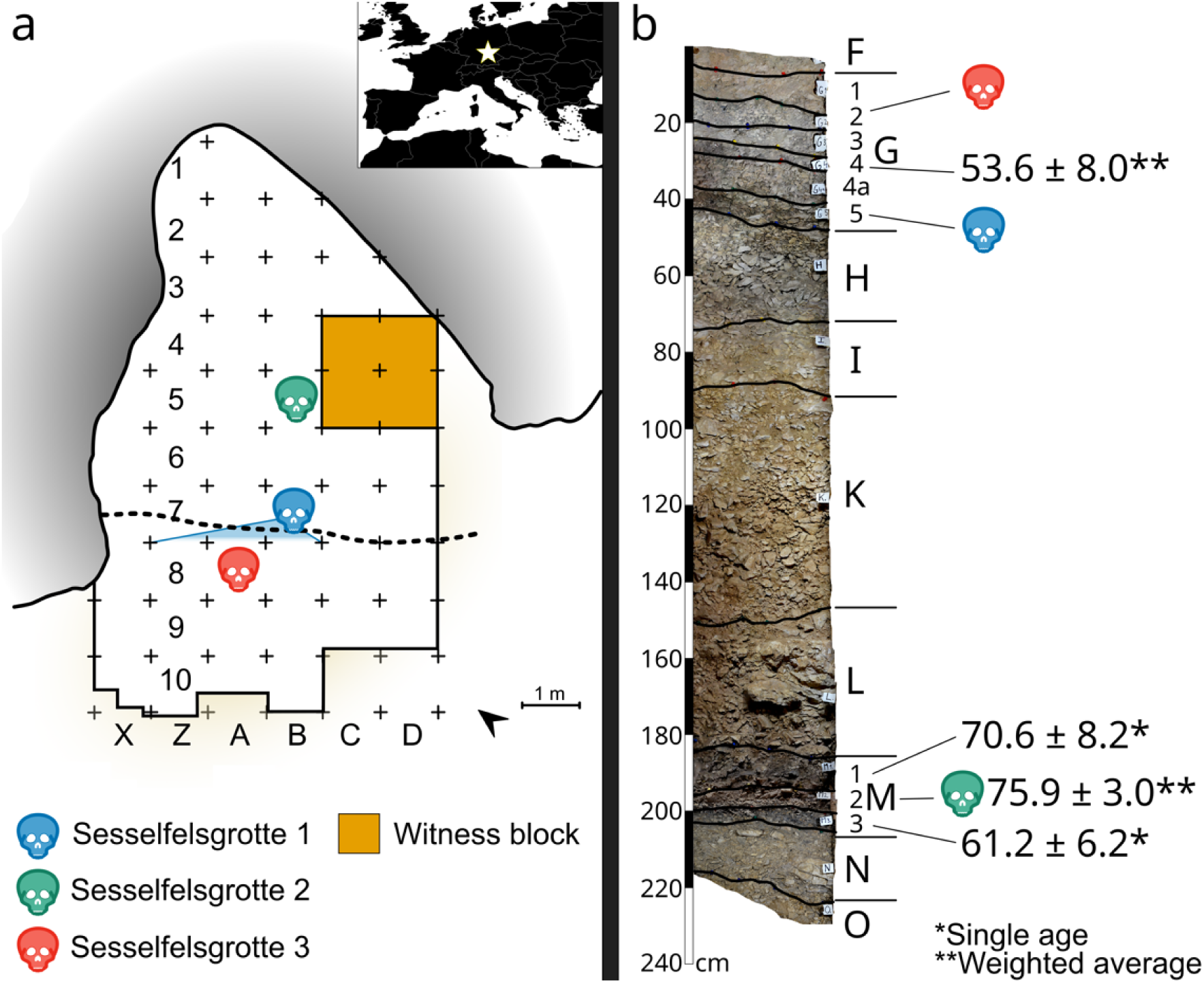
Stratigraphic context of skeletal and dental Neanderthal remains at Sesselfelsgrotte. (a) Planimetry of the cave in its current state, showing the location of the Neanderthal remains Sesselfelsgrotte 1 to 3. The white star in the upper right panel indicates the geographic location of the rockshelter. (b) Stratigraphic log from the witness block in square C5 displaying the vertical position of Neanderthal fossils together with available geochronological data, based on published, but recalculated, thermoluminescence results (Richter et al., 2000). The recalculated thermoluminescence dates exclude samples that Richter (2001) considered as not indicative of Neanderthal occupation.

**Table 1.**
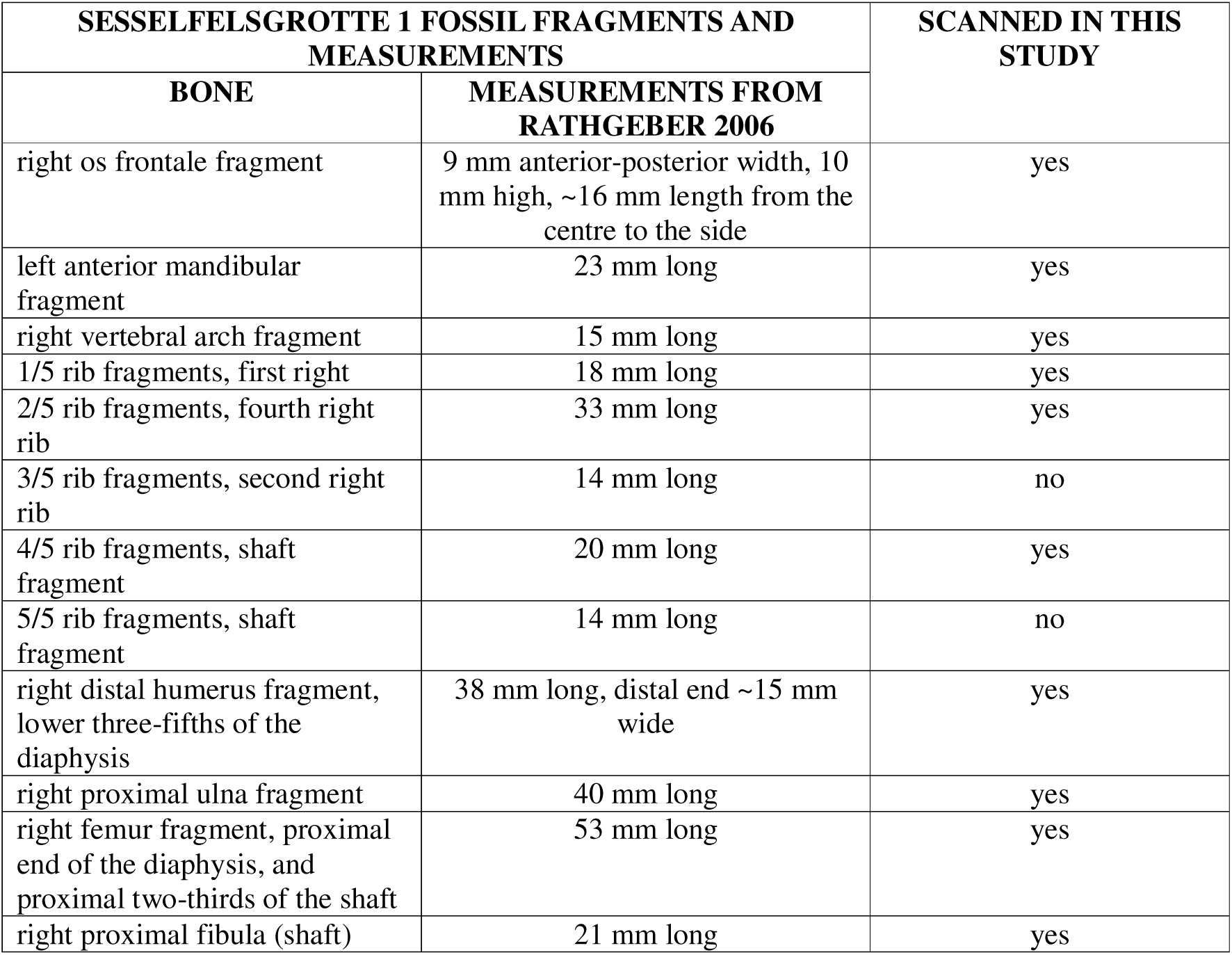
Measurements and anatomical comments on the morphology of bones of Sesselfelsgrotte 1 (extracted from English translation of Rathgeber, 2006 in German). The high fragmentation of the bones precludes conclusive identification of the laterality of the fibula and of the sequence number of the ribs.

| **SESELFELSGROTTE 1 FOSSIL FRAGMENTS AND MEASUREMENTS** |  | **SCANNED IN THIS STUDY** |
| --- | --- | --- |
| **BONE** | **MEASUREMENTS FROM RATHGEBER 2006** |  |
| right os frontale fragment | 9 mm anterior-posterior width, 10 mm high, ~16 mm length from the centre to the side | yes |
| left anterior mandibular fragment | 23 mm long | yes |
| right vertebral arch fragment | 15 mm long | yes |
| 1/5 rib fragments, first right | 18 mm long | yes |
| 2/5 rib fragments, fourth right rib | 33 mm long | yes |
| 3/5 rib fragments, second right rib | 14 mm long | no |
| 4/5 rib fragments, shaft fragment | 20 mm long | yes |
| 5/5 rib fragments, shaft fragment | 14 mm long | no |
| right distal humerus fragment, lower three-fifths of the diaphysis | 38 mm long, distal end ~15 mm wide | yes |
| right proximal ulna fragment | 40 mm long | yes |
| right femur fragment, proximal end of the diaphysis, and proximal two-thirds of the shaft | 53 mm long | yes |
| right proximal fibula (shaft) | 21 mm long | yes |

Rathgeber (2006) noted that almost all Sesselfelsgrotte 1 bones had short length measurements (<2cm, for an overview see Table 1) and appeared highly fragmented (Figure 3). He assigned them to *H. neanderthalensis*, based on the archaeological attribution of the G layers. Moreover, he noted that the Sesselfelsgrotte 1 humerus and femur exhibited distinct robusticity more comparable to *H. neanderthalensis* rather than *H. sapiens*.

**Figure 3:**
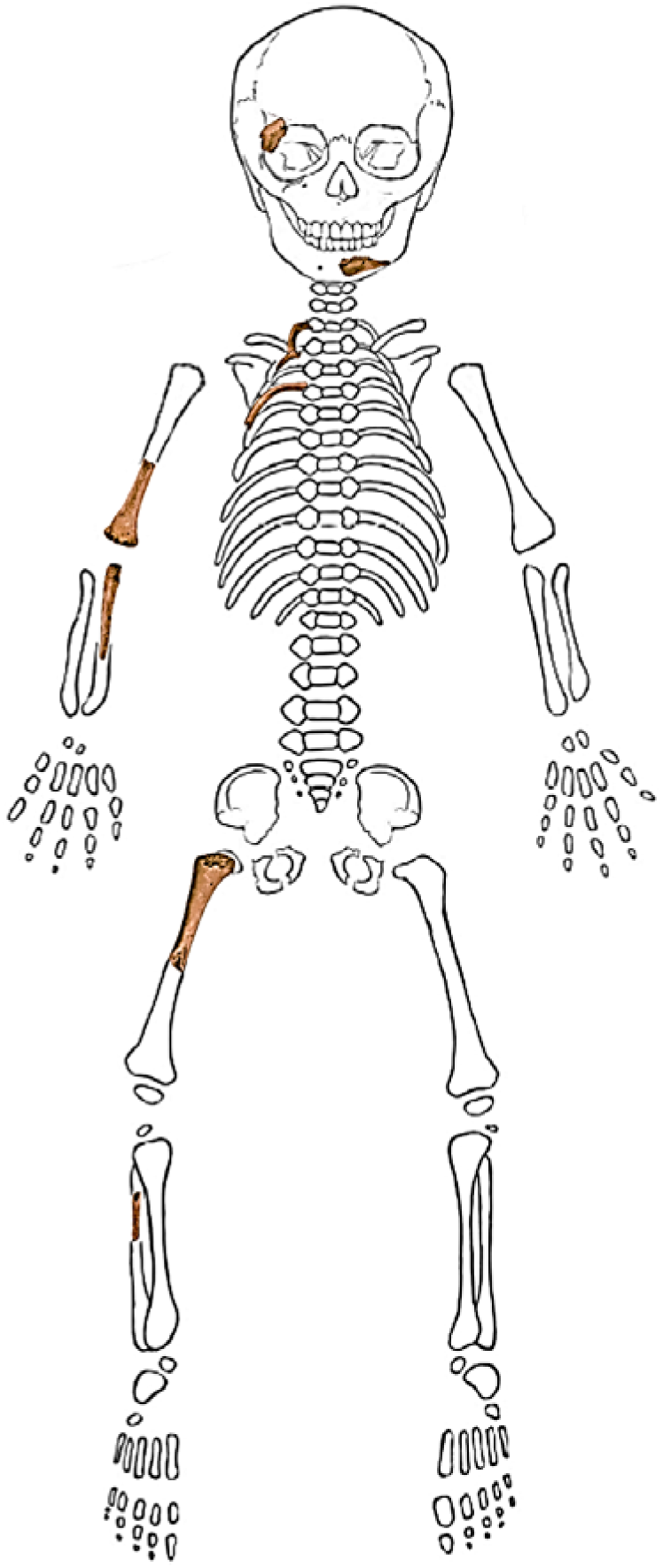
Combined images of the inventory of the fetal remains of Sesselfelsgrotte 1, lacking the bones of the vertebral neural arch because precise identification (i.e., which vertebra) is not conclusive. The high fragmentation of the bones also precludes conclusive identification of the laterality of the fibula and of the sequence number of the ribs.

By comparing his measurements with data from modern human and La Ferrassie foetuses/infants, he inferred these fetal remains to be evidence of a pregnant Neanderthal female who “lost her child in the fetal stage, around 8 months old, during her stay” at the site (translated from German, Rathgeber 2006:53). He acknowledged contention around such interpretations and arguments by others around there potentially being deliberate child burials or simple deposition of Neanderthal remains in cave sediments due to natural processes (Rathgeber, 2006).

New investigations at Sesselfelsgrotte are currently led by A. Barbieri and T. Uthmeier to re-evaluate the site sequence and its assemblages, including the skeletal remains. As part of this project, the femur of Sesselfelsgrotte 1 was sampled for aDNA analysis, revealing that this is indeed a Neanderthal individual (Fotiadou et al., In review). Taphonomic analyses are ongoing to clarify whether these remains were most likely buried as a cadaver by Neanderthals at the site or might have been brought to the shelter by scavengers. In the interim, a preliminary visual taphonomic examination revealed overall preservation suitable for micro-CT analyses in our study with intact internal bone structures. However, most skeletal elements showed chemical alterations of the outer bone surface that may have resulted from digestive processes. Such surface modifications are consistent with partial exposure to gastric acids and suggest that at least some bones might have been affected by carnivore activity. Only in one element, the frontale, did chemical alteration remove a portion of the outer cortex, possibly exposing inner bone structures to further alteration. Previous micro-CT scans of carnivore coprolites suggest that bone fragments ingested by carnivores, when cortical tissue is retained, can preserve intact internal microanatomical structures (Abella et al. 2021). Thus, we expected that most of the Neanderthal remains from Sesselfelsgrotte were suitable to microanatomical study by micro-CT scanning.

### 2.2. Digitization, visualization and analysis

Using micro-CT, we scanned all the skeletal elements of this individual to study their internal bone structure. A Nikon XT H 320 micro-CT scanner with a 0.1 mm thick copper filter at the University of Tübingen was used to digitise the bones with an isotropic spatial resolution of ∼15 to ∼23 microns, 190 kV and 82–90 µA.

Scans were visualized in 3D Slicer (Fedorov et al., 2012) and examined for diagenesis and bone micromorphology. Diagenesis was assessed using the Virtual Histological Index (VHI), which is a virtual complementary method to the Oxford Histological Index (OHI), both of which examine the extent of bioerosion present in bone microstructure (Mandl et al., 2022). This method relies on a visual examination of bone microstructure for preservation that ranges from excellent to poor with corresponding OHI/VHI scores from 5–0 on a 6-point scale progressing from preservation >95% to <5% (Mandl et al., 2022).

The micromorphological bone appearance in the Sesselfelsgrotte 1 individual was examined for qualitative changes in bone tissue arrangement and structure. These were: porosity/vascularity; bone tissue type presence and arrangement (woven, lamellar, primary, secondary, plexiform-like), presence or absence of primary osteons and/or secondary osteons (Goldman et al., 2009; Pfeiffer, 2006; Shapiro & Wu, 2019). We acknowledge that it is essentially not possible to identify woven and lamellar bone collagen orientation from micro-CT scans. Ground histology is appropriate for this type of analysis. Therefore, instead of providing definitive confirmation of woven bone presence, we use the term ‘probably’ to indicate that the bone arrangement seen is not of typical lamellar layering thus pointing to the existence of woven bone tissue. Our micro-CT resolution did not reach cell level details to assess osteocyte lacunae densities, which will remain undescribed here. Our study did not receive permissions to perform destructive analyses, such as ground histology, which would have otherwise offered higher resolution. The fossils could not be removed for scanning at a Synchrotron facility either as we had to limit radiation exposure to enable further sampling for aDNA analysis. With these limitations in mind, we reviewed the entirety of each scanned bone slice by slice, and here provide a summary examination of three main slices per bone taken at three equal segments. As is standard in bone microstructure research, we first describe bone microstructure in each bone on its own and then present overall patterns.

We then compare our findings to published fetal, neonatal, and infant bone microstructure, where available, to check to what extent our observations were consistent with images and data published for various ages (Booth et al., 2016; Caccia et al., 2016; Dhawan et al., 2014; Pfeiffer, 2006; Pitfield et al., 2017; Shapiro & Wu, 2019a; Su et al., 1997). The comparative sample also includes micro-CT scans of the La Ferrassie 4bis and Le Moustier 2 Neanderthals, to which access was granted by the Muséum national d’Histoire naturelle (Paris). Le Moustier 2 originates from a layer which has been dated to 40,3 ±2,6 ka BP by thermoluminescence (Maureille, 2002b; Valladas et al., 1986) and has an estimated age-at-death of <120 days old (Maureille, 2002a) (the right humerus and femur identified as La Ferrassie 4 were originally osteometrically aged as 8,5 fetal months at death by Heim in 1982b, but were later attributed to Le Moustier 2 based on osteometry and tooth calcification [Maureille, 2002a, 2002b]). To the best of our knowledge, there are no dates for La Ferrassie 4bis, nor is the precise stratigraphic origin known, and there are two age-at-death estimates of 12 days post-partum (Heim, 1982b) and 6,5 weeks post-partum (Chevalier et al., 2021). Notwithstanding, dates for La Ferrassie burials have ranged from 54±4 and 43 ka BP based on luminescence (Guérin et al., 2015) to the end of the Middle Paleolithic <52 ka based on more recent combined ^14^C and optically stimulated luminescence (Guérin et al., 2023).

The two deciduous teeth of two other individuals from Sesselfelsgrotte (2 and 3) were first examined macroscopically. Both teeth appear to show evidence of root resorption (but see details below). These specimens were also micro-CT scanned and visualized in 3D Slicer (Fedorov et al., 2012) to examine their internal structures. Hypodense regions found in the dentine were highlighted manually and visualized in 3D to enable examination of their spatial distribution within the teeth.

## 3. Results and Discussion

### 3.1. Bone

All bone slices showed overall very good preservation, which resulted in all VHI scores of >4 meaning preservation >85% (after Mandl et al. 2022; Supplemental Table 1). Only the vertebral specimen was assigned a VHI score of 3.5 due to localized bioerosion which was mostly located in the marrow spaces between bone spicules. We were apprehensive about assigning scores of 5 (>95% preservation) because of the imperfect resolution, but it was clear that any bone diagenesis that was present in the samples was either restricted to the medullary cavity or localized to the sub-periosteum.

All bone fragments examined show bone microstructure consistent with a rapidly forming and growing immature skeleton. We group our descriptions based on developmental pathways: intramembranous ossification in the mandible and the frontal bone fragment, and then endochondral ossification in the vertebral fragment, the rib fragments, and the long bone (humerus, ulna, femur, fibula) fragments. A summary of our observations is provided in Table 2.

**Table 2.** Summary of bone microstructural changes observed in the bones of Sesselfelsgrotte 1.

| Bone markers | Porosity/vascularity | Bone tissue with plexiform-like arrangement | Cortical bone has trabecular-like structure | Bone tissue with primary lamellar arrangement | Woven bone | Primary osteons | Secondary osteons |
| --- | --- | --- | --- | --- | --- | --- | --- |
| **Right frontal bone** |  |  |  |  |  |  |  |
| Slice 1 | High | Yes | Yes | No | Probably | Yes | No |
| Slice 2 | High | Yes | Yes | No | Probably | Yes | No |
| Slice 3 | High | Yes | Yes | No | Probably | Yes | No |
| **Left mandible** |  |  |  |  |  |  |  |
| Slice 1 | High | Yes | Possibly | Possibly | Probably | Yes | No |
| Slice 2 | High | Yes | Yes | Possibly | Probably | Yes | No |
| Slice 3 | High | Yes | Yes | No | Probably | Yes | No |
| **Vertebra** |  |  |  |  |  |  |  |
| Slice 1 | High | Yes | Yes | No | Probably | Yes | No |
| Slice 2 | High | Yes | Partly | Possibly | Probably | Yes | No |
| Slice 3 | High | Yes | Partly | Possibly | Probably | Yes | No |
| **Rib 1** |  |  |  |  |  |  |  |
| Slice 1 | High | Yes | No | No | Probably | Yes | No |
| Slice 2 | High with low region | Yes | No | No | Probably | Yes | No |
| Slice 3 | High with low region | Yes | No | No | Probably | Yes | No |
| **Rib 2** |  |  |  |  |  |  |  |
| Slice 1 | High | Yes | No | No | Probably | Yes | No |
| Slice 2 | High | Yes | No | No | Probably | Yes | No |
| Slice 3 | High with low region | Yes | No | Possibly | Probably | Yes | No |
| **Rib 4** |  |  |  |  |  |  |  |
| Slice 1 | High with low region | Yes | No | No | Probably | Yes | No |
| Slice 2 | High with low region | Yes | No | No | Probably | Yes | No |
| Slice 3 | High with low region | Yes | No | No | Probably | Yes | No |
| **Right humerus** |  |  |  |  |  |  |  |
| Slice 1 | Low | Yes | No | Possibly | Probably | Yes | No |
| Slice 2 | Low with high region | Yes | No | Possibly | Probably | Yes | No |
| Slice 3 | High | Yes | Yes | No | Probably | Yes | No |
| **Right ulna** |  |  |  |  |  |  |  |
| Slice 1 | High | Yes | Yes | No | Probably | Yes | No |
| Slice 2 | High with low region | Yes | No | Possibly | Probably | Yes | No |
| Slice 3 | High with low region | Yes | No | Possibly | Probably | Yes | No |
| **Right femur** |  |  |  |  |  |  |  |
| Slice 1 | High | Yes | Yes | No | Probably | Yes | No |
| Slice 2 | Low | Yes | No | Possibly | Probably | Yes | No |
| Slice 3 | Low | Yes | No | Possibly | Probably | Yes | No |
| **Right fibula** |  |  |  |  |  |  |  |
| Slice 1 | High | Yes | No | No | Probably | Yes | No |
| Slice 2 | High with low regions | Yes | No | Possibly | Probably | Yes | No |
| Slice 3 | High with low regions | Yes | No | Possibly | Probably | Yes | No |

Both the **mandible** and the **frontal** bone fragments (Figure 4a–b), which form through intramembranous ossification, showed slightly different microstructural patterns when compared to the long and other irregular bones that form through endochondral ossification. The shape of the cross-sections is irregular given the predominance of trabecular form over cortical in these bones, but clear high vascularity is still observed in both specimens both on the inferior and superior bone surfaces. Series of primary osteons were clearly seen in the mandible on the superior bone surface, with possible small areas of primary lamellar bone in the same region, but overall, both specimens were still at the plexiform-like stage of formation.

**Figure 4.**
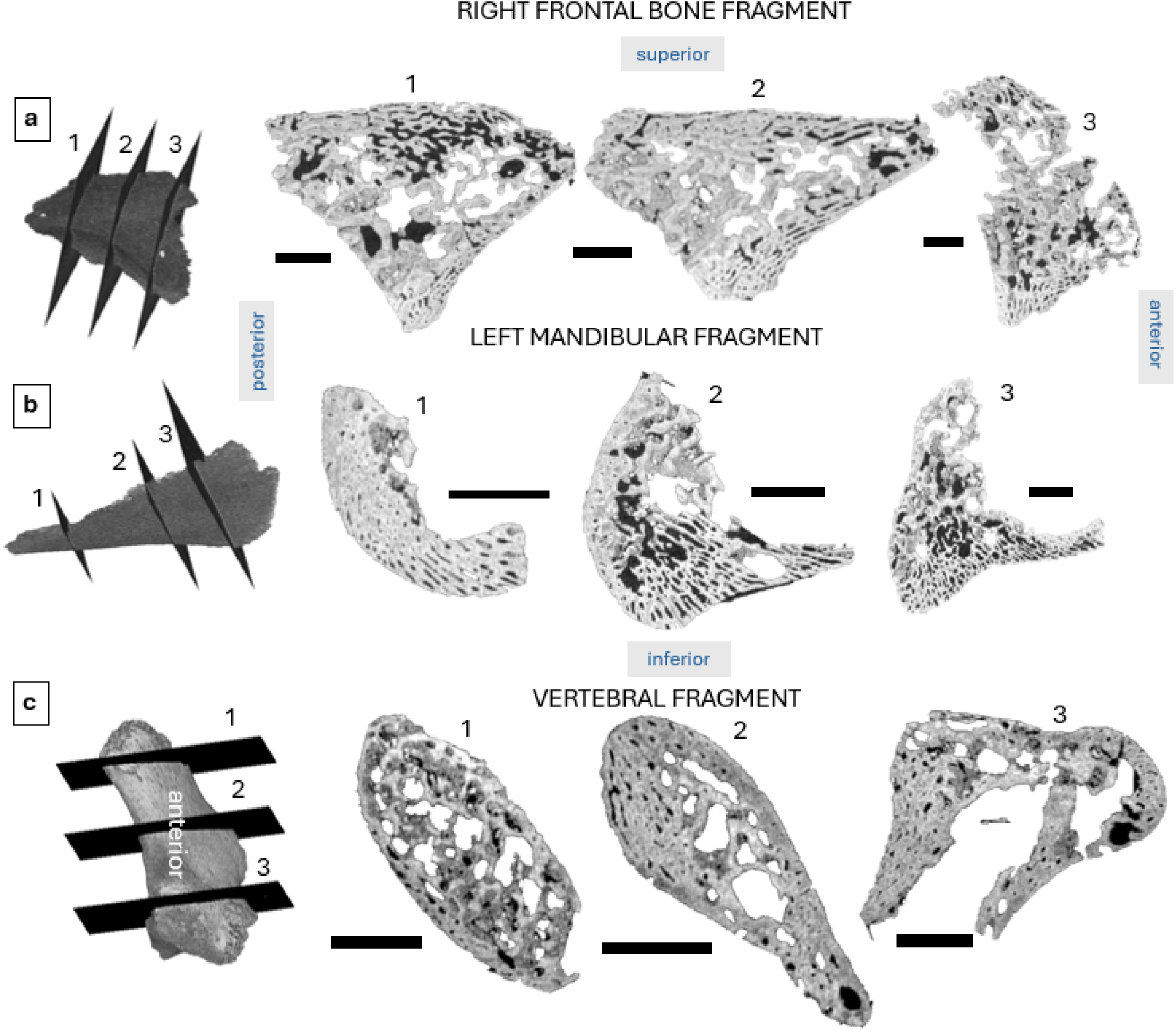
Bone microanatomy visualized in three slices per bone fragment: right frontal bone fragment (a), left mandibular fragment (b), and vertebral fragment (c, most likely anterior view). Scale bars are 10mm.

The **vertebral** fragment (Figure 5c) showed high vascularity throughout its thin cortex, which is consistent of both plexiform-like and trabecular-like structure. There was localized bioerosion covering the trabeculae within the internal spongy component of the vertebra, which obscures the microanatomy, but the cortical area exhibited possible small areas of primary lamellar bone. The bone tissue in the three **ribs** (Figure 5) shared similar micro-morphology with high vascular densities arranged in a plexiform manner and isolated primary osteons. The medullary cavity here was small in proportion to the cortical area with remnants of trabecular bone. Rib 2 showed a possible small region of primary lamellar bone on the cutaneous surface, but this was only noticeable in the rib region located towards the sternal end (see slice 3) and not in the other parts or the other ribs (see slices 1–2).

**Figure 5.**
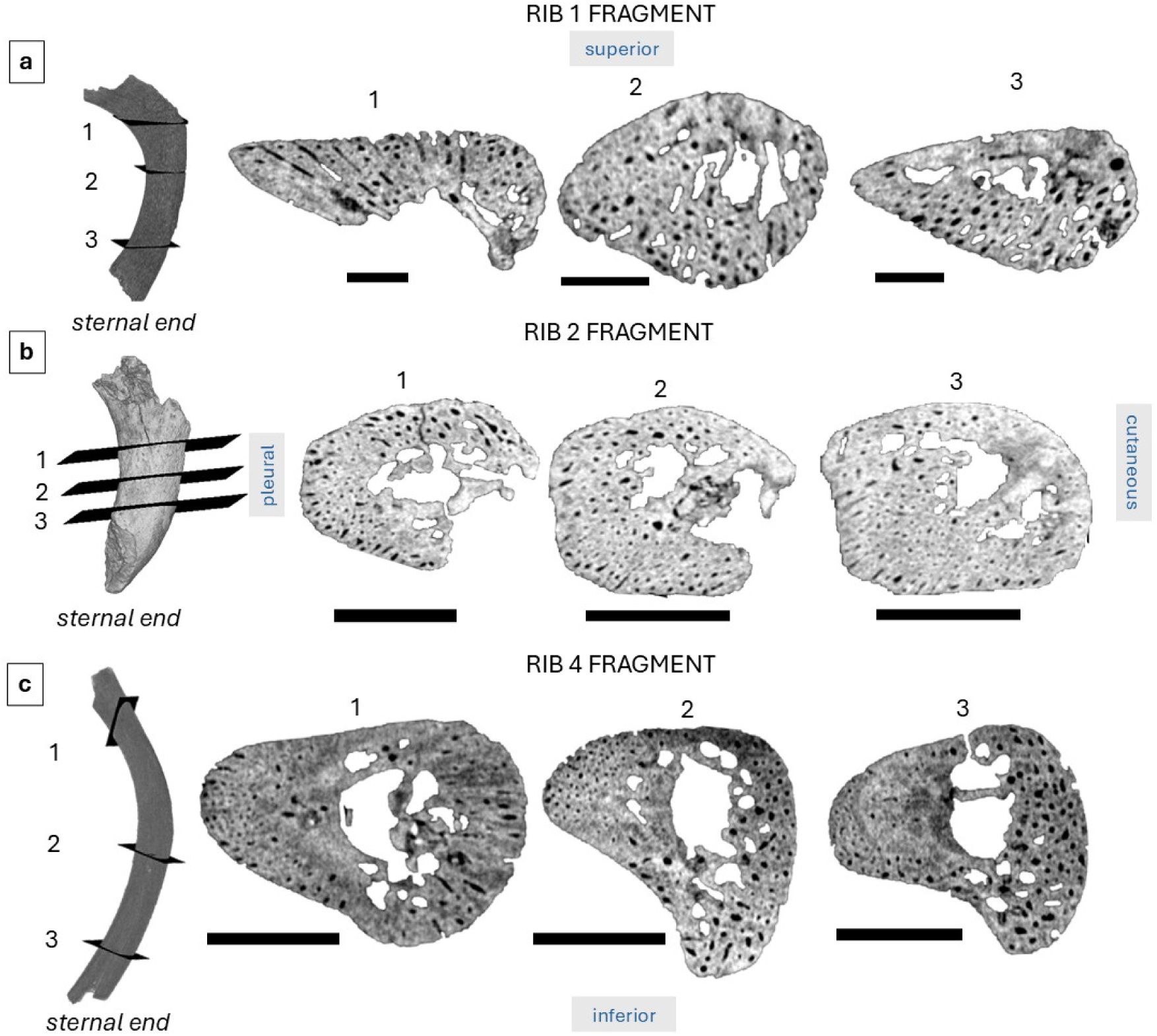
Bone microanatomy visualized in three slices per rib fragment. Scale bars in panel a are 5 mm, while panels b–c show scale bars that measure 10 mm.

The long bone shafts (Figure 6) all shared very similar micro-morphology, with the largest of the four bones (femur and humerus) showing overall higher density of compact bone on the intra-cortical and sub-periosteal region than the ulna and the fibula. This was particularly evident on the slices viewed in the midshaft area of these bones. The **humerus** (Figure 6a) fragment near the midshaft showed distinct isolated trabeculae and a proportion of bone tissue: medullary cavity nearing 1:1 ratio in terms of occupied space. It showed the lowest porosity (in terms of vascular spaces) compared to femur and the ulna. There were primary osteons seen along with the remnants of plexiform-like woven bone complex spanning the entirety of the compact wall. There were also possible regions of primary lamellar bone nearer the midshaft on the poster-medial aspect. Out of all the long bones, the **ulna** (Figure 6b) showed the highest vascularity of a combination of radial and transversely oriented canals of the plexiform-like pattern. At the shaft, its medullary cavity did not retain trabecular form and the compact bone layers showed only sporadic primary osteons restricted to the endosteal regions of bone, but with possible areas of primary lamellar bone on the postero-lateral aspect. Nearing the proximal end where spongy bone appeared, there were clear trabeculae present surrounded by highly vascularized thin cortex.

**Figure 6.**
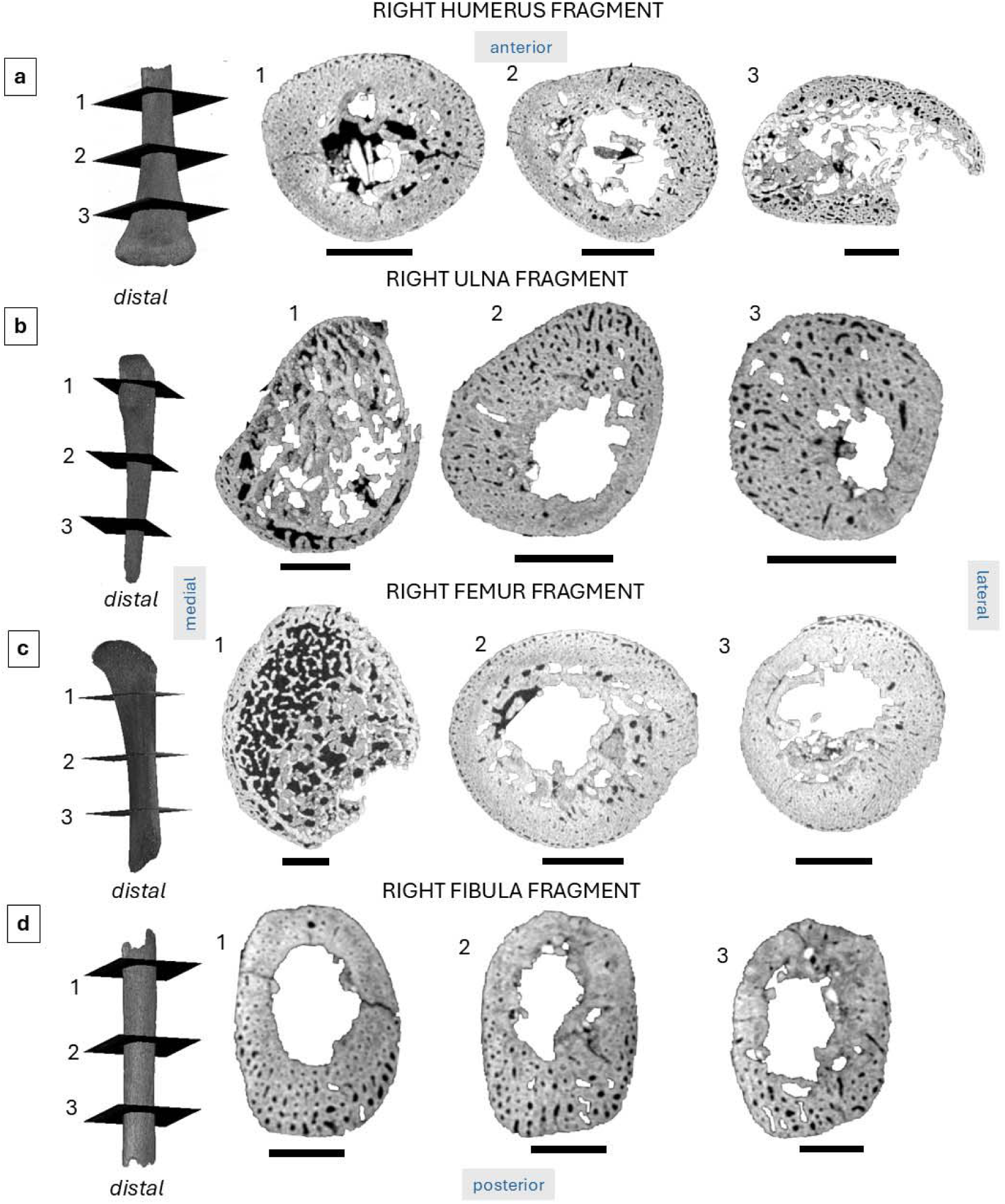
Bone microanatomy visualized in three slices per bone fragment: right humerus fragment (a, posterior view), right ulna fragment (b, posterior view), right femur fragment (c, posterior view), and right fibula fragment (d, posterior view). Scale bars in panels a–c are 10mm, while panel b shows scale bars of 5mm length.

The **femur** (Figure 6c) showed the largest cross-section compared to the other long bones with emptier medullary space than that of the humerus with a limited number of trabeculae restricted to the endosteal region. Vascularity characterized by plexiform-like arrangement was obvious on the sub-periosteal region, particularly on the antero-medial surfaces, but it transitioned into primary osteon lamellar form intra-cortically and towards the endosteal areas with localized radial canals. The shaft of the **fibula** (Figure 6d) showed an oval cross-section with a well-defined medullary cavity retaining only a small number of endosteally peripheral trabeculae. The cortical bone was well vascularized with primary osteons and small regions of primary lamellar bone on the antero-lateral surfaces, and areas of plexiform-like structure on the poster-medial aspect.

Once compared (summary in Table 3) to the information and images available in the literature and anatomical sources (Booth et al., 2016; Sawada et al., 2004; Shapiro & Wu, 2019a; Su et al., 1997), and scans for the <6,5 weeks post-partum La Ferrassie 4bis and <120 days old Le Moustier 2 (Figure 7), the highest similarity in the microanatomical appearance was with the femoral midshaft micro-CT slices from Romano-British archaeological pre-term individuals (Booth et al., 2016) aged 30–36 weeks (some shown in Figure 1e–h). Comparisons with femoral and humeral shaft micro-CT slices from La Ferrassie 4bis and Le Moustier 2 did not quite agree, with Le Moustier 2 showing stronger compaction of the cortical walls, while La Ferrassie 4bis showing intense vascularity. Sesselfelsgrotte 1 scans had a mix of these features. However, in La Ferrassie 4bis, the femur and humerus were highly impacted by bioerosion so it is difficult to comment further on vascularity structures. The Le Moustier 2 femur and humerus also showed some bioerosion in the cortical bone, obscuring microanatomical structures, but the compactness of bone and well defined cortical and trabecular spaces indicated a relatively older individual than Sesselfelsgrotte 1.

**Table 3.** Summary of our observations consistent with information and images available in the literature.

| **Bone/specimen** | **Reference** | **Age** | **Sesselfelsgrötte 1 comparisons**<br>(younger/older/similar) |  |  |
| --- | --- | --- | --- | --- | --- |
| **Bone** | **Ref** | **Age** | **Rib** | **Humerus** | **Femur** |
| Modern human fetal femur midshaft histology | Su et al. 1997 | 14–26 weeks | - | - | older |
| Modern human fetal femur microanatomy (IS6) | Booth et al. 2016 | 34–36 weeks | - | - | **similar** |
| Modern human fetal femur microanatomy (IS7) | Booth et al. 2016 | 30–32 weeks | - | - | **similar, potentially slightly older** |
| Modern human fetal femur microanatomy (IS10) | Booth et al. 2016 | 30–32 weeks | - | - | **similar, potentially slightly older** |
| Neanderthal femur midshaft histology | Sawada et al. 2004 | ~2 years old | - | - | younger |
| Modern human femur midshaft histology | Sawada et al. 2004 | ~1 year old | - | - | younger |
| Modern human femoral and humeral fetal parallel-fibred lamellar bone deposited on fetal woven bone cores | Shapiro & Wu 2019 | ~4–4.5 gestational months | - | older | older |
| Modern human juvenile rib | Pfeiffer, 2006 | 0–1 yrs | younger | - | - |
| Modern human juvenile humerus | Pitfield et al. 2017 | 0–1.9 yrs | - | younger | - |
| Modern human fetal femur | Dhawan et al. 2014 | 21+5 weeks | - | - | older |
| Modern human neonatal femur | Caccia et al. 2016 | <1 year | - | - | younger |
| La Ferrassie 4bis | Figure 7 | <6,5 postnatal weeks | - | older | some similarities |
| Le Moustier 2 | Figure 7 | <120 days | - | younger | younger |

**Figure 7.**
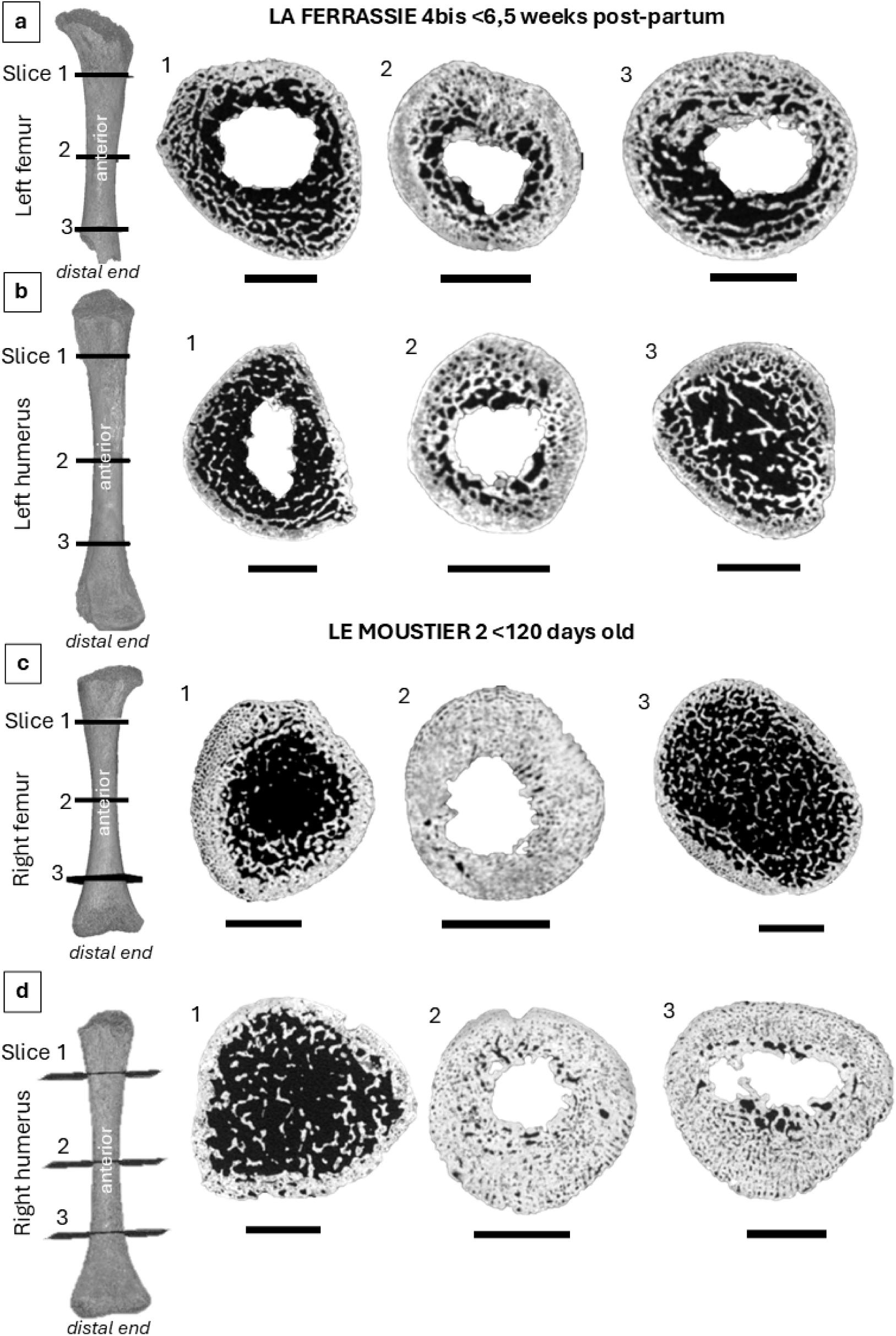
Bone microanatomy visualized in the micro-CT scans of the humerus and the femur in La Ferrassie 4bis (a–b) and Le Moustier 2 (c–d). Scale bars are 10mm.

Overall, the micro-CT scans of bone microanatomy of fragments of the femur, humerus, ulna, fibula, three ribs, mandible, vertebra, and frontal bone of Sesselfelsgrotte 1 revealed skeletal tissue structure partly consistent with late third trimester gestation in *H. sapiens*, but showing regions of advanced bone formation that is typically seen in modern human neonates. Our findings support the age estimates based on macroscopic measurements presented by Rathgeber (2006) around the 8^th^ fetal month mark. Our analysis also brings new information about Neanderthal *in utero* development of the skeleton, showing remarkable similarities to modern human fetal bone microstructure this late in the final trimester of pregnancy. This supports what has been previously proposed using Neanderthal and modern human long bone microstructure around both taxa sharing similar developmental infant trajectories up until about 1–2 years of age after which modern humans depart from Neanderthal type bone microstructure (Sawada et al., 2004).

Our examination of bone microstructure agreed with fetal bone formation and growth in modern humans. The very characteristic plexiform-like bone structures and probable woven bone seen in all the Sesselfelsgrotte 1 bones indicate clearly that they belonged to an individual whose skeleton had not yet been exposed to significant mechanical and temporal factors that would otherwise stimulate the formation of more mature bone tissues. We refer to this tissue type as plexiform-like because it is thought that modern humans do not deposit plexiform tissues as defined in other larger mammals that have a rapid juvenile growth (Goldman et al., 2009; Pfeiffer, 2006). In those mammals, plexiform bone takes a distinct brick-wall pattern of superimposed layers of bone. In modern human fetal bone, this tissue is more disorganized with irregular vascular spaces which can orient themselves in many directions. The presence of this bone tissue in combination with primary osteons is evidence that bone is experiencing rapid growth, which is expected for foetuses. While plexiform-like bone tissue has been previously documented in older modern human children who undergo growth spurts (Martin & Burr, 1989), the macroscopic measurements and comparative analyses undertaken by Rathgeber (2006) would rule out a much older child. Further, none of the Sesselfelsgrotte 1 bones showed any obvious primary lamellar bone arranged in clear successive layers to begin forming circumferential lamellar bone, both of which would indicate the slowing down of bone formation and deposition of bone that is better organized than woven and plexiform-like. Sawada et al. (2004) noted a lack of circumferential lamellae in the Dederiyeh 1 Neanderthal child at 1.5–1.7 years, but there was presence of secondary and primary osteons. We identified primary osteons, but no secondary osteons.

The overall complex of bone tissues in our specimens was clearly a combination of woven, plexiform-like, and some primary osteons, with regions of the humerus and femur showing progression towards increased compactness and more structured organization. These areas may indicate a slight advancement in bone growth that would be consistent with conclusions made by Sawada et al. (2004) that Neanderthal cortical bone histology (at least in the femur) has features different from modern humans despite similarities in primary bone in the first 1–2 years. If that is the case, we of course cannot confirm the underlying causes of possible bone growth advancement in the Sesselfelsgrotte 1 Neanderthal. Neanderthal skeletal robusticity both at the juvenile and adult stages has been consistently linked to climate adaptation, lifestyles of various physical demands, varied diets, and faster development than that of modern humans (Churchill, 2014; Pomeroy, 2023). Considering the subject of our study are perinatal bones, we can only propose an alternative or a combination of factors such as dietary, maternal, and genetic, since it is known that these factors can determine the speed with which the fetal skeleton forms (Goodfellow et al., 2010). While the biomechanics of a Neanderthal foetus are essentially impossible to study, the existing body of knowledge about modern human fetal movement (e.g., reviewed in Nowlan, 2015) and morphological similarities between Neanderthals and modern humans, suggest that Neanderthals would have experienced similar ranges of fetal movement yielding *in utero* mechanical stimulation (e.g., twitching; stretching; full body, limb, head/neck/jaw movements) supporting bone formation. Possible areas of the beginning of compactness in the femur and the humerus could point to these fetal movements being correspondingly strong to Neanderthal robusticity (Nowlan et al., 2010).

Our observations of bone microstructure are limited by the imperfect resolution of the micro-CT scans, which most obviously hindered our ability to evaluate cell-level information at the osteocyte lacuna level, but we avoided destructive and radiation analyses due to their destructive nature. These other methodological approaches could have aided in gathering more data, particularly clarifying any more localized bioerosion and allowing us to conduct quantitative analyses, such as estimating primary osteon counts and densities. The fragmentary preservation of the fossils also means we cannot comment on bone formation in other bones or the remaining parts of bones. Another large limiting factor in our study is the bioerosion seen in the comparative Neanderthal fetal bone microanatomy. This means all of our conclusions are based on comparisons with modern humans and limited information for Neanderthals, such as ground histology of the Dederiyeh 1 child (Sawada et al., 2004) and imperfect microanatomical micro-CT scans (Figure 9). Deciduous tooth crowns could have also confirmed whether this Neanderthal was in fact born and estimate age-at-death by using the neonatal line and enamel incremental lines (Macchiarelli et al., 2006), but regrettably there are no teeth preserved in Sesselfelsgrotte 1. Future research in this area should aim to undertake multi-methodological approaches at higher resolution to further contribute to our current understanding of Neanderthal fetal bone microstructure.

### 3.2. Deciduous teeth

The root of the Sesselfelsgrotte 2 tooth is almost fully resorbed (Figures 8–9), while the Sesselfelsgrotte 3 tooth is very incomplete, with only about ¼ of the crown preserved and the root also almost completely absent (Figures 10–11). The anatomical identification as a deciduous lower left second molar is, thus, tentative. Visual inspection of the root appears to show resorption of ¾ of its length (Figure 10)**Figure**, but it is unclear if it may be broken and not resorbed.

**Figure 8.**
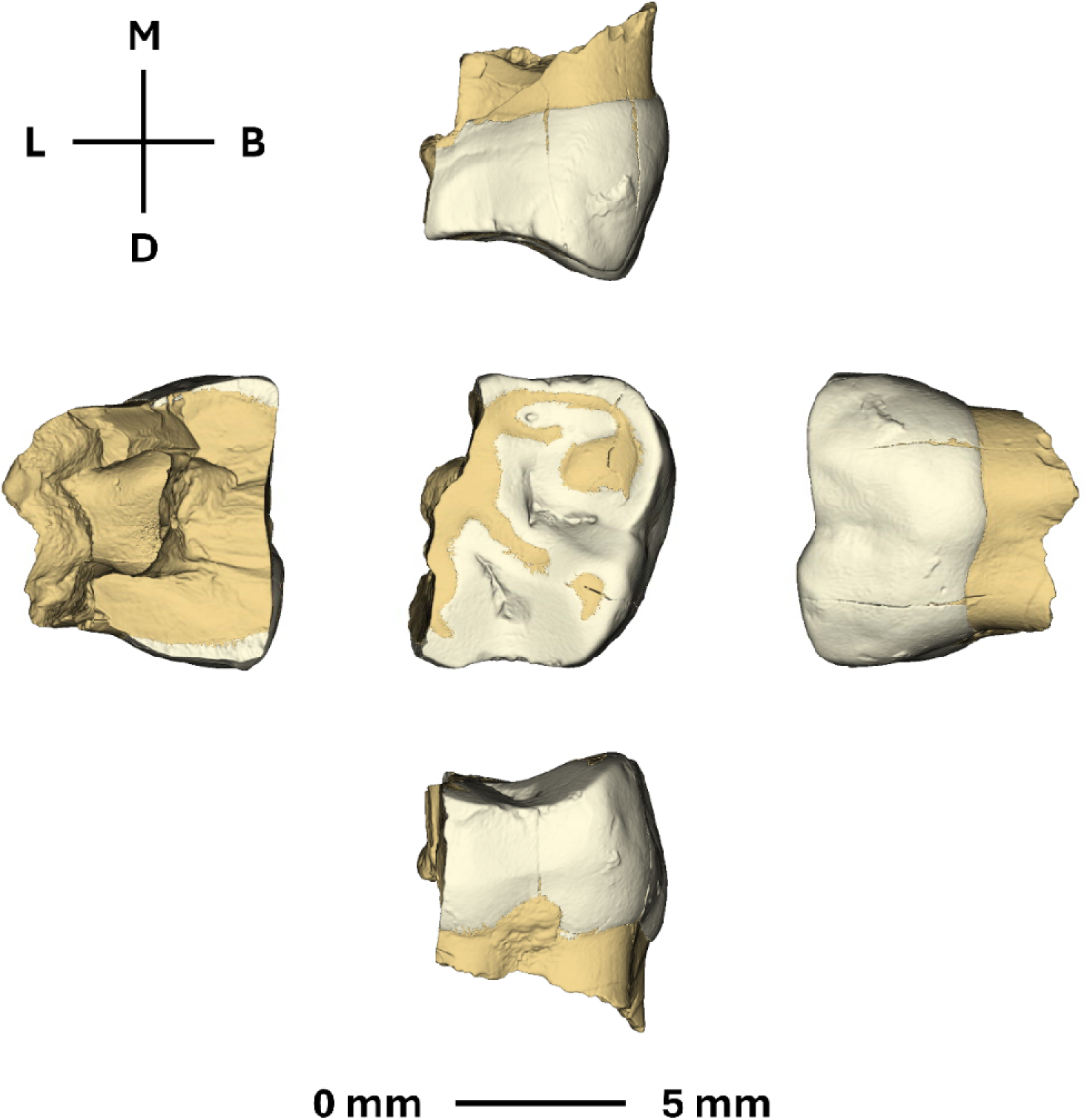
Sesselfelsgrotte 2 (upper left second deciduous molar).

**Figure 9.**
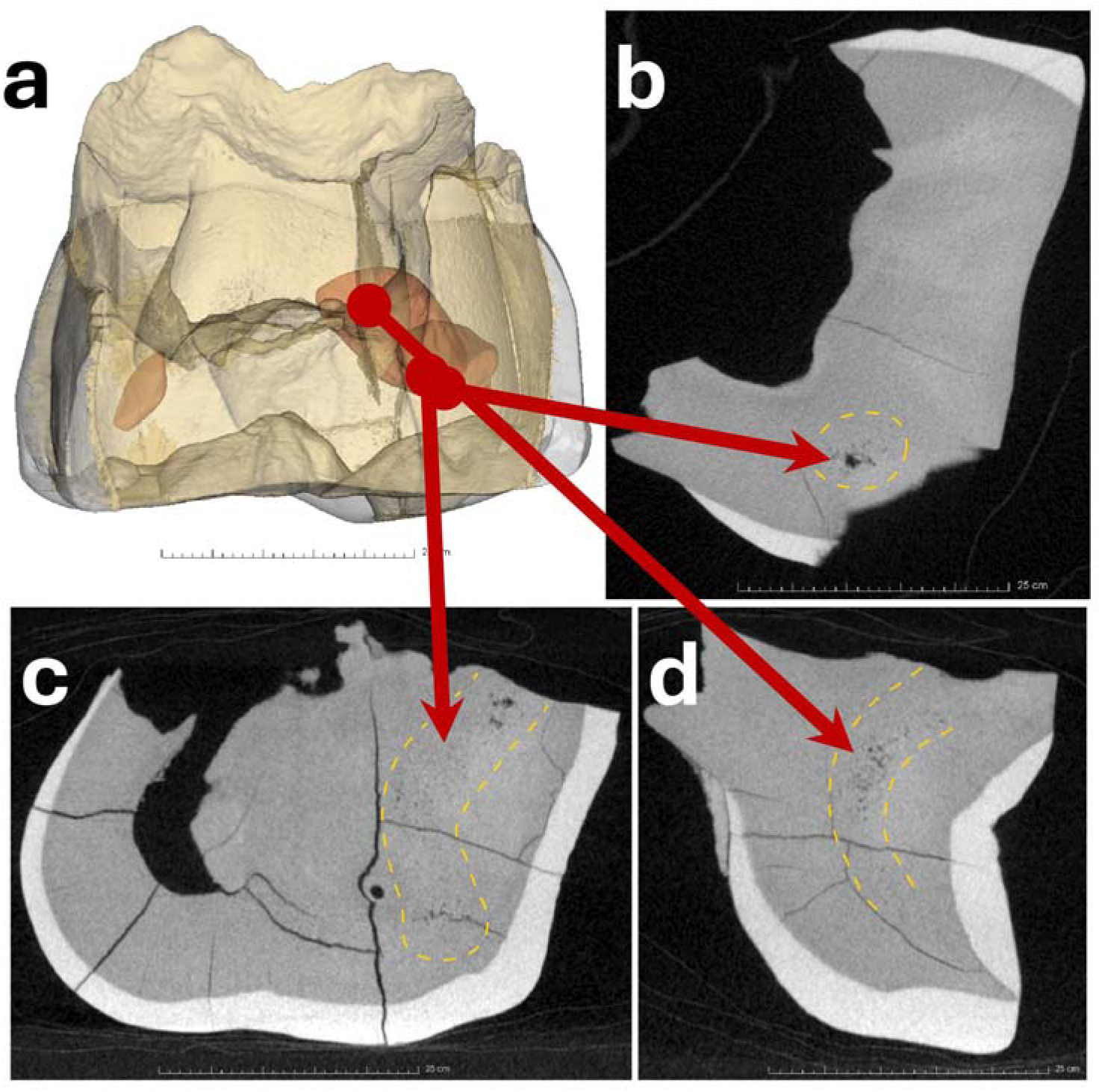
Sesselfelsgrotte 2: (a) Spatial distribution and examples of (b, c) hole-like or (c, d) pinpoint hypodensities in the crown circumpulpal dentine in the upper second deciduous molar from Sesselfelsgrotte. d) shows a region where these hypodensities are clustered.

**Figure 10.**
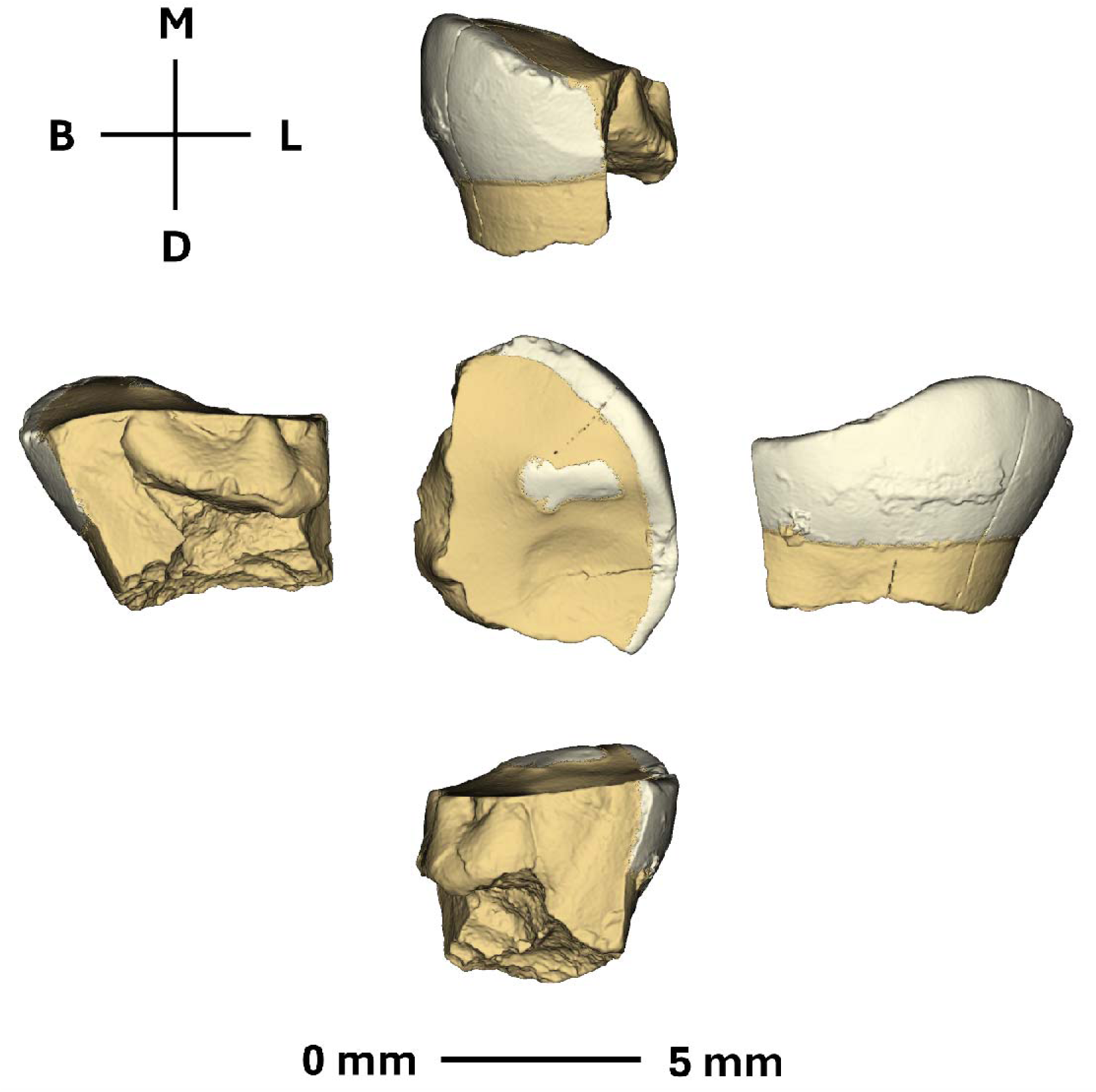
Sessefelsgrotte 3 (lower left second deciduous molar).

**Figure 11.**
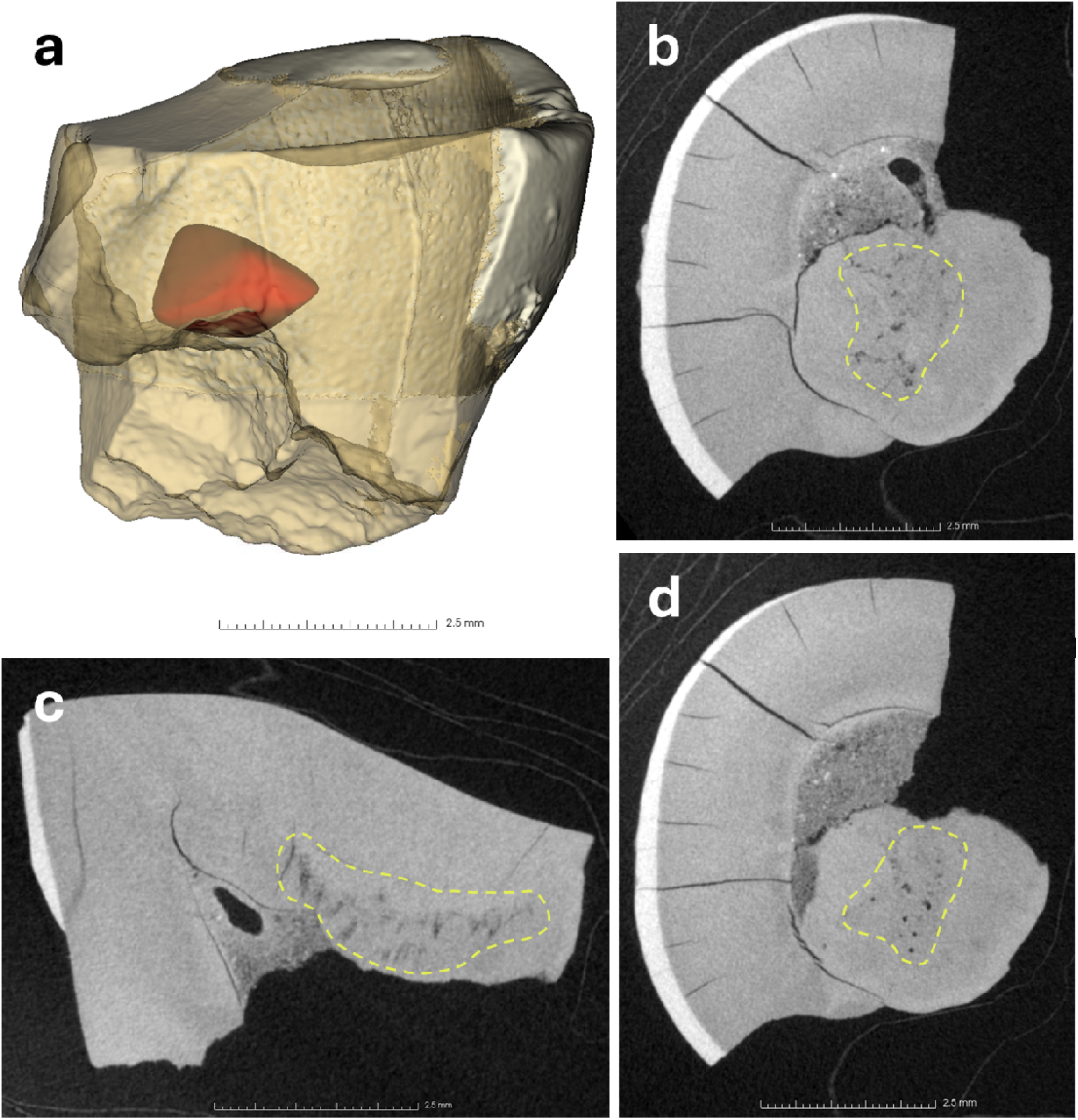
(A) Spatial distribution and (b–d) examples of hypodensities in the crown circumpulpal dentine in Sesselfelsgrotte 3.

Although the Sessefelsgrotte 2 tooth is not fully preserved (part of the lingual half is absent), the tooth crown is worn with dentine exposure in the mesial-buccal cusp, less so in the distal-buccal cusp and, based on the preserved region, presumably more in the lingual half of the tooth (approximately dental wear stage 5 of the wear scale of Smith, 1984). This, together with root resorption, precludes observation of the whole dentinal record. Several hypodensities (areas of less mineralization) were detected in the primary coronal circumpulpal dentine. Specifically, three *foci* of reduced density were detected in the mesial and distal regions of the circumpulpal dentine away from the enamel dentine junction (and so from the mantle dentine in the crown; Figure 9a). These regions are visible as clusters of dark-grey/black pinpoints and/or larger black hole-like regions (Figure 9b–d). Although the tooth presents multiple internal fractures, these hypodensities are not present in the affected regions.

Sesselfelsgrotte 3 is incomplete, with extensive occlusal dental wear and a small enamel island remaining on the preserved occlusal surface (which suggests an approximate dental wear stage 6–7 in the scale of Smith, 1984, if this regions is representative of the full occlusal surface). Similar to Sesselfelsgrotte 2, several hypodensities were detected within the primary coronal circumpulpal dentine. Specifically, a region of reduced density was detected in the circumpulpal dentine and distant from the enamel dentine junction (and so from the mantle dentine in the crown; Figure a) in which dark-grey/black pinpoints and/or larger black hole-like regions are found. Although the tooth presents multiple internal fractures, these hypodensities are not associated with the affected regions.

The hypodensities detected in both Sesselfelsgrotte 2 and 3 are consistent with the appearance of documented IGD in historical modern human teeth imaged with micro-CT (Colombo et al., 2019; Veselka et al., 2019). It is impossible to definitively ascribe an underlying aetiology to IGD in fossil remains. The presence of IGD has been clinically associated with inborn errors of vitamin D metabolism (Karemore et al., 2022; Seow et al., 1989). Additionally, IGD has been noted in modern human archaeological individuals with macroscopic skeletal evidence of rickets or osteomalacia, likely due to nutritional vitamin D deficiency when the environmental context is considered (D’Ortenzio et al., 2016; Snoddy et al., 2024; Veselka et al., 2019). However, any disorder that inhibits availability of calcium for the mineralization of hard tissues will produce a similar suite of skeletal lesions (rickets/osteomalacia) and this is likely true for IGD as well (Vautour & Goltzman, 2018; Vlok et al., 2023).

Because pathological IGD arises due to a systemic disturbance in pre-dentine mineralization, it occurs in both sides of the teeth in band-like formations along incremental growth lines and across several teeth (D’Ortenzio et al., 2018). There is some debate over how much IGD needs to be present in a region to be classified as “pathological” as opposed to IGD that forms from the normal retraction of odontoblasts within dentine tubules (developmental IGD). However, developmental IGD should be present in the region most proximal to the dentine-enamel junction, rather than deeper in the primary dentine, and is not expected to form in discrete bands along incremental growth lines (D’Ortenzio et al., 2016; Nanci, 1994). Additionally, comparison of ground histological sections and micro-CT data found that IGD typically scored as developmental is too small and widely spaced to be visualized on micro-CT scans (Colombo et al., 2019). Thus, the IGD in Sesselfelsgrotte 2 and 3 is likely abnormal and suggestive of a period of poor mineral metabolism during growth. Potential causes include nutritional vitamin D deficiency and/or dietary calcium deficiency, a period of poor calcium absorption due to intestinal disease, or temporary renal compromise (Bikle et al., 2019; Vautour & Goltzman, 2018). Because the severity and number of episodes of IGD in x-linked hypophosphatemic rickets is variable and we only have a partial development record, we cannot exclude this as a possibility (Boukpessi et al., 2006).

Due to a lack of visibility of dentine incremental lines, it is not possible to identify the chronological age that the IGD occurred but based upon the duration of Neanderthal deciduous molar formation it is possible to suggest a range of biological ages when the poorly mineralized dentin may have formed. Upper second molar enamel of modern humans starts to form around three months before birth (Mahoney et al., 2025). Dentine secretion is initially slightly in advance, and the crown is formed approximately 13 to 17 months after birth (Mahoney et al., 2025; Sunderland et al., 1987). The Krapina Neanderthal deciduous first molar crowns formed over a slightly shorter postnatal period (Mahoney et al., 2021), which was likely due to the slightly accelerated rates of formation that have been reported for some Neanderthal deciduous (Macchiarelli et al., 2006) and permanent teeth (Ramirez Rozzi & Bermudez de Castro, 2004; Smith et al., 2010). Assuming that the dm2 of Sesselfelsgrotte 2 and 3 formed over a slightly shorter, or a period that was equivalent to modern human dm2’s, then the IGD would occur somewhere between the third trimester and up to the middle of the second postnatal year. Most likely the period of poor mineralization occurred towards the middle and end of this period given the position of the IGD in Figures 9 and 11.

Previous studies suggest that several aspects of Neanderthal femoral curvature and non-adult skull morphology were pathologically induced, and specifically related to rickets/osteomalacia (Czarnetzki, 2000; Ivanhoe, 1970; Ivanhoe & Trinkaus, 1983; Klaatsch, 1901). Despite those early interpretations, such Neanderthal morphological aspects have been dismissed as related to rickets/osteomalacia (De Groote, 2011; Ivanhoe & Trinkaus, 1983), and currently the earliest reported hypothesized cases of vitamin D deficiency originate from layer B of the Tabun cave, from which other dental remains have been dated to ∼82–92 kya (Coppa et al., 2005). Specifically, one adolescent first molar and one left mandibular first molar from an individual with 6–12 years of age at death is said to show clear evidence of interglobular dentine (Sognnaes, 1956). Additionally, one maxillary left central incisor from a young adolescent displayed some granularity but no large regions of interglobular dentine (Sognnaes, 1956). Although these teeth are hypothesized to be Neandertal (Sognnaes, 1956), this tentative classification lacks confirmation (Brickley et al., 2017). Moreover, D’Ortenzio et al. (2018) argue that these cases are probably of developmental origin, rather than induced by vitamin D deficiency. More recently, one undisputedly Neanderthal tooth from Lakonis, Greece, dated to 38–44 Ka (Harvati et al., 2003; Panagopoulou et al., 2002), was also reported to display interglobular dentine (Smith et al., 2009). Thus, presently, reports of interglobular dentine in Neanderthals are restricted to the Lakonis individual (Smith et al., 2009) and to the tentative cases from Tabun (Sognnaes, 1956). The Sesselfelsgrotte 2 and 3 deciduous Neanderthal teeth add to these few reported possible vitamin D deficiency induced IGD cases and are one of the earliest, if not the earliest. However, as mentioned above, IGD is a clinically common finding in modern humans and an evidenced-based threshold for the levels of serum vitamin D at which IGD begins to form have not yet been established. Vitamin D deficiency is a major health issue in modern times and the frequency at which it is found in clinical thin sections may or may not be related to the global prevalence of this condition. Work to explore the relationship between serum vitamin D status and IGD formation in modern human teeth is ongoing.

## 4. Conclusions

In this study we examined bone microanatomy in the fossil bones of Sesselfelsgrotte 1 Neanderthal previously estimated to have died as a perinate, based on osteometry. Using non-invasive micro-CT, we found that all bones showed microanatomical patterns consistent with modern human fetal growth in the final trimester of pregnancy approaching 8–9 months, which agrees with the prior macroscopic measurements. We observed a combination of immature, probably woven, plexiform-like, and isolated primary osteonal bone tissues which are typical for rapidly growing and fast forming skeletons. This is consistent with a younger Neanderthal than, for example, the 1.5–1.7-year-old Dederiyeh child bone histology specimens which had secondary osteons. In the context of what is currently known for Neanderthal fetal and infant growth, our study contributes a new perspective that their fetal growth might have had slight acceleration in long bone formation but overall is consistent with modern humans until at least 1–2 years. Additionally, we have observed the potential earliest, approximately 75,000 ya, and clearest presence of interglobular dentine that is likely pathological in a Neanderthal lineage. Although we refrain from attributing a definitive cause for this lesion, it is indicative of poor mineral metabolism and is the earliest evidence of metabolic bone disease in a non-anatomically modern human to date.

## Acknowledgements

We thank Christina Kyriakouli and Gabriel Ferreira from Senckenberg Centre for Human Evolution and Palaeoenvironment for operating the micro-CT scanner at the University of Tübingen, Agnes Fatz for high resolution photographs of the skeletal and dental elements from Sesselfelsgrotte, Thomas Higham and Andreas Pfemeter for discussions about preliminary ^14^C dating results, and the Muséum National d’histoire Naturelle (Paris) for facilitating access to the micro-CT scans of La Ferrassie 4bis and Le Moustier 2.

## Ethics statements

Ethics clearance was not applicable. Access to all fossils was granted by curating institutions.

## Data availability statement

The fossils are permanently stored and curated at the Prehistoric and Protohistoric Museum of the UFG Institutes based at the Friedrich-Alexander-Universität (FAU) in Erlangen (Bavaria). The fossils are not currently displayed, to see the materials please contact Thorsten Uthmeier. The micro-CT scans of all specimens examined in this study will be made available open access from Morphosource upon publication of this article.

## Author contributions

JJM: first draft, study design, methods, data analysis and interpretation; RMG: study design, data analysis and interpretation, methods, materials, software, data curation; AMS: methods, data analysis and interpretation; KP: methods, data analysis and interpretation; FD: methods, materials, data analysis and interpretation; PM: methods, data analysis and interpretation, TR: methods, materials, data analysis and interpretation; CP: materials, methods, software, data interpretation, funding; TU: materials, methods, software, data interpretation, funding; AB: study design, materials, methods, software, data interpretation, data curation, funding. All authors edited the manuscript.

