## Supplemental Table 1 for "Early development of Neanderthals revealed through virtual microanatomy"

***Equal first authors**

**^a^Corresponding authors:**

Justyna J. Miszkiewicz, School of Social Science, University of Queensland, 4067 Brisbane, Australia

Ricardo M. Godinho, Interdisciplinary Centre for Archaeology and the Evolution of Human Behaviour (ICArEHB), Universidade do Algarve, 8005-139 Faro, Portugal

Alvise Barbieri, Interdisciplinary Centre for Archaeology and the Evolution of Human Behaviour (ICArEHB), Universidade do Algarve, 8005-139 Faro, Portugal

**Supplemental Table 1.** Virtual Histological Index scores assessing virtually visible bioerosion, measured on a 0-5 scale with 0 indicating poor preservation and 5 indicating perfect preservation.

| **Bone** | **MicroCT Slice 1** | **MicroCT Slice 2** | **MicroCT Slice 3** |
| --- | --- | --- | --- |
| right os frontale | 4 | 4 | 4 |
| left anterior mandible | 4 | 4 | 4 |
| right vertebral arch | 3.5 | 3.5 | 3.5 |
| 1/5 rib fragments, first right | 4 | 4 | 4 |
| 2/5 rib fragments, fourth right rib | 4 | 4 | 4 |
| 3/5 rib fragments, second right rib | n/a | | |
| 4/5 rib fragments, shaft fragment | 4.5 | 4.5 | 4.5 |
| 5/5 rib fragments, shaft fragment | n/a | | |
| right distal humerus fragment, lower three-fifths of the diaphysis | 4.5 | 4.5 | 4.5 |
| right proximal ulna fragment | 4 | 4 | 4 |
| right femur fragment, proximal end of the diaphysis, and proximal two-thirds of the shaft | 4.5 | 4.5 | 4.5 |
| right proximal fibula (shaft) | 4 | 4 | 4 |
